# OptiFlex: video-based animal pose estimation using deep learning enhanced by optical flow

**DOI:** 10.1101/2020.04.04.025494

**Authors:** XiaoLe Liu, Si-yang Yu, Nico Flierman, Sebastian Loyola, Maarten Kamermans, Tycho M. Hoogland, Chris I. De Zeeuw

**Affiliations:** Faculty of Mathematics, University of Waterloo, Waterloo, Canada; Dept. of Neuroscience, Erasmus MC, Rotterdam, The Netherlands; Netherlands Institute for Neuroscience, Royal Academy of Arts and Sciences, Amsterdam, The Netherlands

**Author notes:** These authors contributed equally (XiaoLe Liu, Si-yang Yu).

## Abstract

Deep learning based animal pose estimation tools have greatly improved animal behaviour quantification. However, those tools all make predictions on individual video frames and do not account for variability of animal body shape in their model designs. Here, we introduce the first video-based animal pose estimation architecture, referred to as OptiFlex, which integrates a flexible base model to account for variability in animal body shape with an optical flow model to incorporate temporal context from nearby video frames. This approach can be combined with multi-view information, generating prediction enhancement using all four dimensions (3D space and time). To evaluate OptiFlex, we adopted datasets of four different lab animal species (mouse, fruit fly, zebrafish, and monkey) and proposed a more intuitive evaluation metric - percentage of correct key points (aPCK). Our evaluations show that OptiFlex provides the best prediction accuracy amongst current deep learning based tools, and that it can be readily applied to analyse a wide range of behaviours.

## Introduction

Being able to precisely describe and quantify animal behaviour has profound implications in fields such as neuroscience or psychology. Whereas the human visual system can readily interpret raw video data on animal behaviour at a qualitative level, precise quantification of body positions and movements requires complex computational analyses. The major challenge is to reliably extract essential information of animal behaviour for downstream tasks, such as motion clustering^1, 2^ or sensorimotor correlations^3–6^, to allow for quantitative descriptions. This problem can be addressed by articulated animal pose estimation, which consistently tracks predetermined key points on a given animal.

Though key point tracking can be done with great accuracy through human labelling on a frame-by-frame basis, it usually incurs considerable time and labour cost, limiting the annotation of large datasets. The need for accurate, fast and scalable tracking of animal behaviours has therefore driven several efforts to automate animal pose estimation using both marker-based and markerless tracking. Marker-based tracking of key points usually involves placing reflective markers that can be detected with a camera system^7, 8^. Alternatively, one can use accelerometers to directly readout movement acceleration^9^. Marker-based tracking has the advantage of providing straightforward processing of object location and animal pose^10^. However, its invasive nature can disrupt animal behaviour.

Markerless tracking on the other hand circumvents the stress and workload associated with marker placement and could be the method of choice, if it matches the accuracy of marker-based tracking. Early markerless tracking used Kinect cameras^11^ or multi-camera systems^11^, setting constraints on the simplicity and versatility of the experimental setting in which animal behaviour can be measured. Developments in deep learning based computer vision techniques, especially convolutional neural networks, provide critical building blocks to further advance markerless tracking^12–15^. Accordingly, advances in deep learning based human pose estimation have inspired novel tools in animal pose estimation. For example, DeepLabCut^16^ is based on the feature detector from DeeperCut^17^, and StackedDenseNet from DeepPoseKit^18^ is a variation on Fully Convolutional DenseNets^19, 20^ that are stacked in the fashion of Stacked Hourglass^21^.

While these approaches have brought meaningful advances to animal pose estimation^12^, they directly transferred computer vision techniques developed for tracking humans to lab animals, omitting key differences in shape and size. With regard to pose estimation, the most important differences concern the number and size of key points. Indeed, the number of key points can heavily influence the size and complexity of pose estimation models and the ratio between key point sizes affects the accuracy and meaning of the evaluation metric.

Since humans have very similar body shapes, human pose estimation datasets usually contain an identical number of key points, and a constant ratio between key point sizes. As a consequence, pose estimation models capture the same amount of complexity and evaluation metrics can use the size of a specific joint as a threshold. For instance, one common metric in human pose estimation is that a key point estimation is correct if its distance from the ground truth is less than half the size of the head (PCKh@0.5)^21–23^. Lab animals, on the other hand, vary greatly in shape and size. This means that while certain datasets can be readily modelled by small and simple models, others require more complex models with a larger number of parameters. Such variability carries the risk of overfitting when using a complex model on a simple dataset, and underfitting when using a simple model to fit a complex dataset. Our work here presents a flexible base model and an adaptive, yet intuitive, evaluation metric - percentage of correct key points (aPCK), to account for such variability. We adopted datasets of four different lab animals species, i.e. mouse, fruit fly, monkey, and zebrafish, to entrain and test our models under various natural experimental conditions.

Given that animal behaviours are usually recorded in video format, we employ the first video-based model architecture. Most animal pose estimation models to date^16, 18, 24^ focus on predicting key points using a single video frame. By taking an image-based approach, these models neglect the sequential nature of these images, and thus ignore valuable temporal context. In contrast, our video-based approach utilizes a short sequence of images, comprising a target frame and several adjacent frames, to make a key point prediction. With a video-based approach such as OptiFlex features from various time points can all be used at the same time to enhance prediction accuracy and robustness against temporary obstruction of key points.

## Results

Our overall workflow consists of a video-based model architecture along with a graphical user interface (GUI) for training data annotation and augmentation (**Fig. 1a**). The model architecture is comprised of a base model and an optical flow model. The base model makes initial predictions on the input images, and the optical flow model converges the predictions into a single prediction for the target frame of the input images (**Fig. 1b**). Details regarding GUI can be found in Supplemental Material and each component of the model architecture will be discussed in detail in the following sections.

**Fig. 1.**
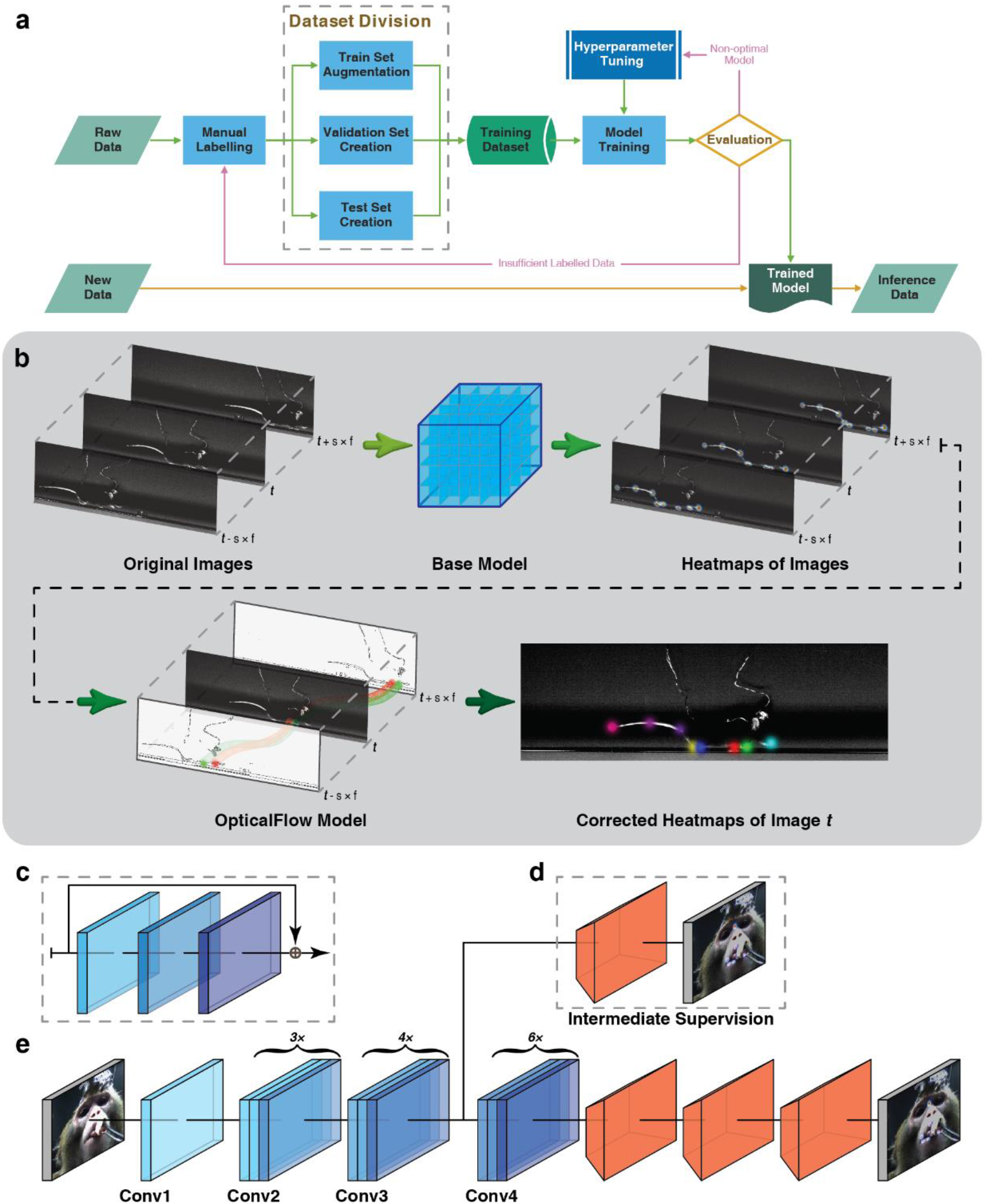
Workflow and model architecture of OptiFlex. **a**, Overall data preprocessing and model training pipeline. **b**, Given skip ratio and frame range *f*. For a target frame with index *t*, we first gather a sequence of 2*f* + 1 images with index from *t* − *s* × *f* to *t* + *s* × *f*. The base model makes a prediction on each of the images to create a sequence of heatmap tensors with index from *t* − *s* × *f to t* + *s* × *f*. The OpticalFlow model takes the entire sequence of heatmap tensors and outputs the final heatmap prediction for index *t*. **c**, Diagram of a “bottleneck” building block commonly used in ResNet backbone, consisting of 3 convolutional layers and a skip connection. **d**, Optional intermediate supervision for FlexibleBaseline through additional loss calculation between heatmap label and intermediate results from ResNet backbone after a single transposed convolution represented by orange trapezoidal blocks. **e**, Standard FlexibleBaseline with intermediate supervision after Conv3 block. Note that each Conv block consists of multiple “bottleneck” building blockings (taken from panel **c**).

### Percentage of Correct Key Points (aPCK) vs. Root Mean Square Error (RMSE)

RMSE is currently the default evaluation metric in recent work on animal pose estimation^16, 18, 24^. While RMSE can be useful as the loss function for neural networks, this metric does not intuitively reflect prediction quality and can sometimes be misleading.

RMSE can be faulty both when comparing the performance of different models using the same dataset and when comparing the performance of the same model applied to different datasets. When comparing two models using the same dataset, the model with a slightly larger RMSE is not necessarily worse. This is because animal key points can be a few pixels in size; therefore, prediction values that differ by a few pixels can both be correct. Moreover, since all points of a given RMSE from ground truth form a circle around the ground truth location and an acceptable region for a key point can be of any shape, points with the same RMSE from ground truth can both be valid or incorrect predictions (**Fig. 2**). When comparing the same model using different datasets, RMSE can be even worse. A model with a given RMSE can make perfect predictions in a dataset with larger joint sizes, while being completely inaccurate in another dataset with smaller joint sizes (**Fig. 2**). All these subtle flaws of RMSE can eventually be amplified by the aforementioned variability that is innate to animal datasets. This combination of biases may explain why RMSE is not often used to evaluate human pose estimation^21–23^.

**Fig. 2.**
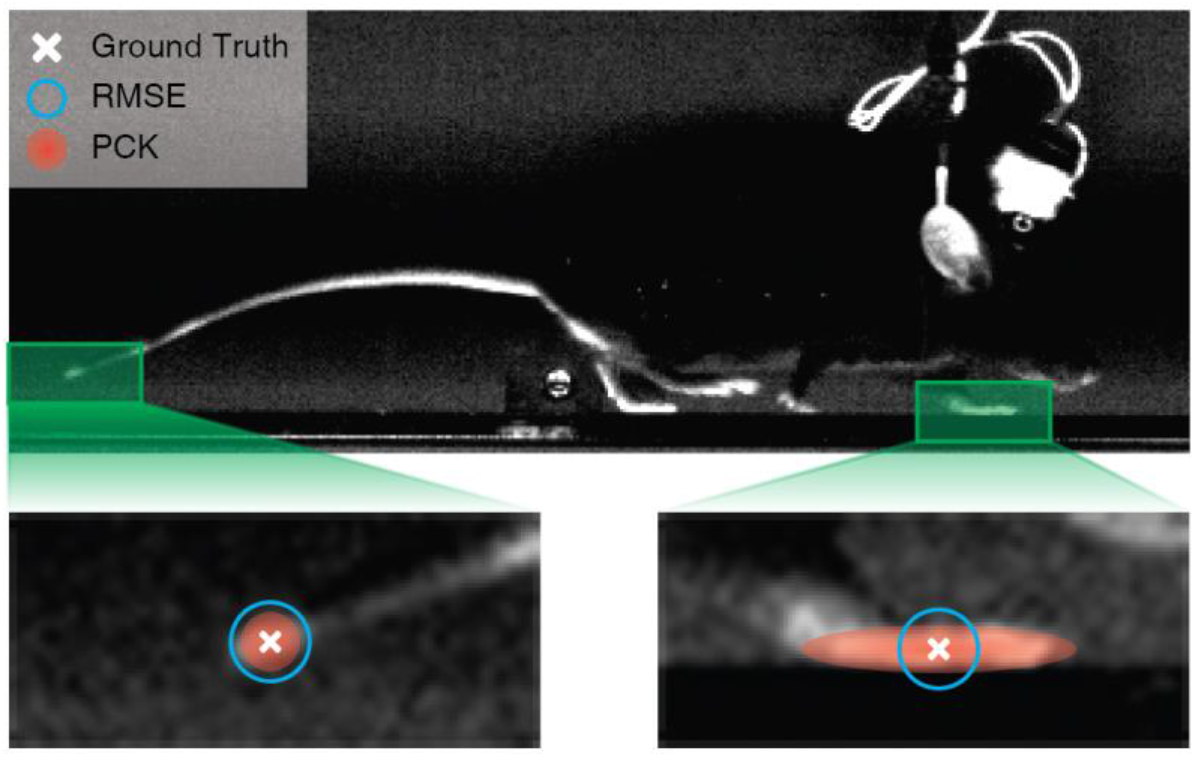
Comparison of evaluation methods. The ground truth is defined by a human generated label. Points with a given RMSE forms a circle around the ground truth. The heatmap label, used by aPCK, can be defined by the human labeller to cover the entire ROI, but the RMSE circle can be too large and include points that are not on the paw or it can be too small and miss valid points of the paw.

A more revealing metric for model performance is the percentage of key point predictions that land in an accepted region around the ground truth. To avoid confusion with the predefined notion of PCK^25^ in computer vision, we denote the percentage of correct key points for animals as aPCK. The aPCK accounts for the variability of different animal datasets by defining the area that is considered acceptable or “correct” with the key point heatmap labels as generated by human subjects. The heatmap generated for each joint follows a truncated 2D Gaussian distribution with peaks at the human labelled points. The heatmap primarily functions to distinguish points closer to the ground truth, as they have higher weight than points further away even when all points in the heatmap are considered “correct”. Normalization to account for various scales of the animal and image will naturally happen in the human labelling process as users can specify label size. For the current study we define the prediction as “correct”, if the predicted key point location has a readout value > 0 of the ground truth label heatmap. This makes the metric very flexible, giving the user control over acceptable regions. Hence, we use aPCK as the default metric for all following evaluations.

### FlexibleBaseline Structure of OptiFlex

In principle, any imaged-based model that predicts a set of heatmaps of key point likelihood can be used as a base model. For OptiFlex, we devised a base model, FlexibleBaseline, that can serve as a baseline for future work in animal pose estimation. FlexibleBaseline’s overall structure consists of a ResNet^26^ backbone, 3 transposed convolution layers, and a final output layer (**Fig. 1c-e**). The ResNet backbone is a section of the original ResNet that uses weights pre-trained on ImageNet^27^ and can output after any of the Conv blocks from ResNet. It also allows optional intermediate supervision anywhere between the input layer and the backbone output layer, usually after a Conv block. The different options for backbone output and intermediate supervision endow the model with ample flexibility. The 3 transposed convolution layers all have filters of size 13×13 (i.e. the window for scanning through the input image or intermediate tensors is 13×13) with strides depending on output location of ResNet backbone and with a modifiable number of filters in each layer depending on dataset complexity. The final output layer always has the same number of filters as the number of prediction key points, each with filter size 1×1 and stride 1. This structure is inspired by the Simple Baselines^28^, which attained state-of-the-art results in many human pose estimation challenges in COCO 2018^29, 30^.

### FlexibleBaseline Performance and Comparison Against Previous Models

A number of pose estimation models were included in our comparison: LEAP, DeepLabCut, and DeepPoseKit (StackedDenseNet). For each animal, the labelled datasets were divided according to the dataset division table (**Table S2**). All models were trained using a training set, hyperparameters were selected using a validation set, and final evaluation was done on a test set. For all datasets, our FlexibleBaseline achieved the best performance amongst all of the models in terms of prediction error rate (**Fig. 3**); in particular, FlexibleBaseline significantly outperformed all other models on mouse side view and zebrafish datasets. For the monkey and fruit fly datasets, FlexibleBaseline also had the lowest error rates, but these differences were not significant, because for these two datasets the other models were also performing virtually perfectly. A detailed comparison of model accuracy is provided in Supplemental Material **Table S5** and a video comparison of tracking results is shown in **Video S1-a~d**.

**Fig. 3.**
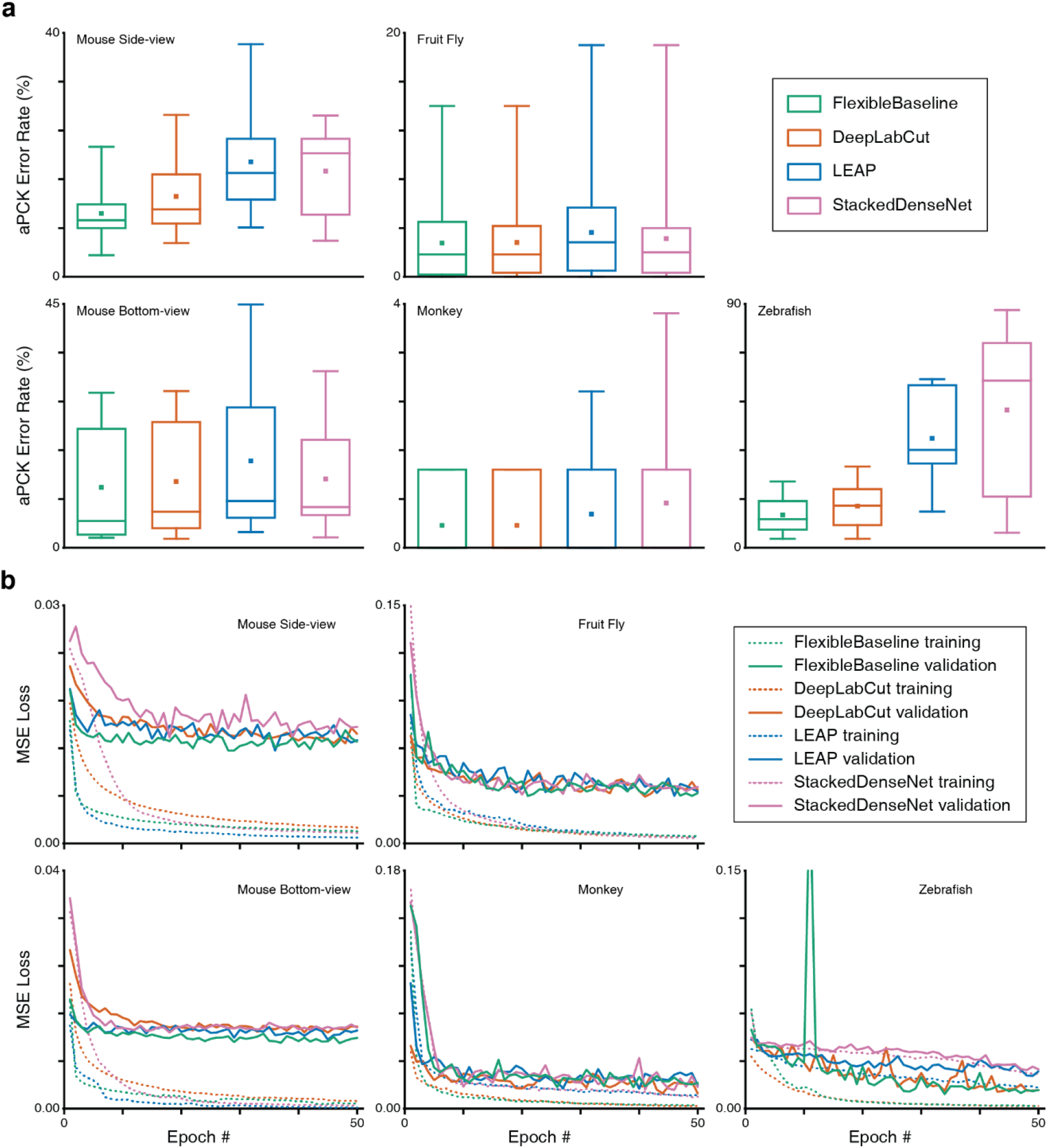
Comparison of model error rates and loss curves on different datasets. **a**, Test set prediction error rates represented in box plots with minimum, first quartile, median, third quartile, and maximum. The small square in the middle of each box plot represents the mean error rate. Note that some monkey results do not show whiskers (parameters of variability), because of the virtually perfect predictions. **b**, MSE training and validation loss; note the peak in zebrafish reflects the fact that the optimizer landed in a poor spot early in the training process.

The inference speed of FlexibleBaseline was measured as the time the model takes to predict heatmaps from preprocessed input tensors of a particular dataset. The measurements were done on VM instances of identical configuration on Google Cloud (see Computing Environment). To account for potential variability, the same prediction process was run 16 times, and the final results reflect the averages of these runtimes. For real-time inference with a batch size of 1, FlexibleBaseline had an average per image inference speed of 35ms for the fruit fly test set, 18ms for the monkey test set, 25ms for the zebrafish test set, 12ms for the mouse bottom view test set, and 14ms for the mouse side view test set. For larger batch sizes, inference speed can still increase. For example, with a batch size of 128, FlexibleBaseline has an average per image inference speed of 26ms for the fruit fly test set, and with a batch size of 256, FlexibleBaseline has an average per image inference speed of 16ms for the monkey test set, 24ms for the zebrafish test set, 6ms for the mouse bottom view test set, and 8ms for the mouse side view test set.

### Flexibility in Resource Constrained Situations

The flexibility of FlexibleBaseline derives from the fact that a user can select output from any of the 5 Conv blocks from ResNet50 and specify the number of filters in the last three transpose convolution layers. Different combinations of output block and filter numbers can vary greatly in the number of parameters, and thereby training and inference speed, while retaining a comparable accuracy across datasets. This flexibility gives users a very favourable trade-off between speed and accuracy when necessary.

In fact, smaller models may outperform larger models in resource constrained situations, where not enough labelled data are available or the hardware does not support a large number of epochs. To simulate these conditions, we tested FlexibleBaseline with 3 different hyperparameter settings using a significantly reduced number of training steps and a minimal amount of annotation on the mouse side view and fruit fly datasets. The number of parameters in these models decreased from more than 25 million in the standard version to less than 2 million in the small version (see Methods for detailed model and training setup, and Supplemental Material for performance of the 3 versions under the previous non-constrained training setup).

Recent animal pose estimation models^18, 24^ suggest that reasonable accuracy can be achieved with as few as 100 labelled frames on the fruit fly dataset. We thus started with only hundreds of frames for both datasets. Independent from the versions of FlexibleBaseline, training with 100 frames in the fruit fly dataset yielded prediction error rates that were comparable to those obtained with the full dataset (**Fig. 4a**). When we gradually reduced the number of labelled frames, we observed a natural increase in prediction error rates with all versions. The standard version of FlexibleBaseline had the lowest error rate on the vast majority of the tested datasets. In case of the mouse side view dataset, the models had significantly higher prediction error rates with 300 labelled frames, so we explored training with a geometrically increased number of labelled frames. Our results indicated that under resource constrained training setups, small and reduced versions outperformed the standard version in all of the tested mouse side view datasets, while prediction error rates showed a plateau beyond 600 labelled frames.

**Fig. 4.**
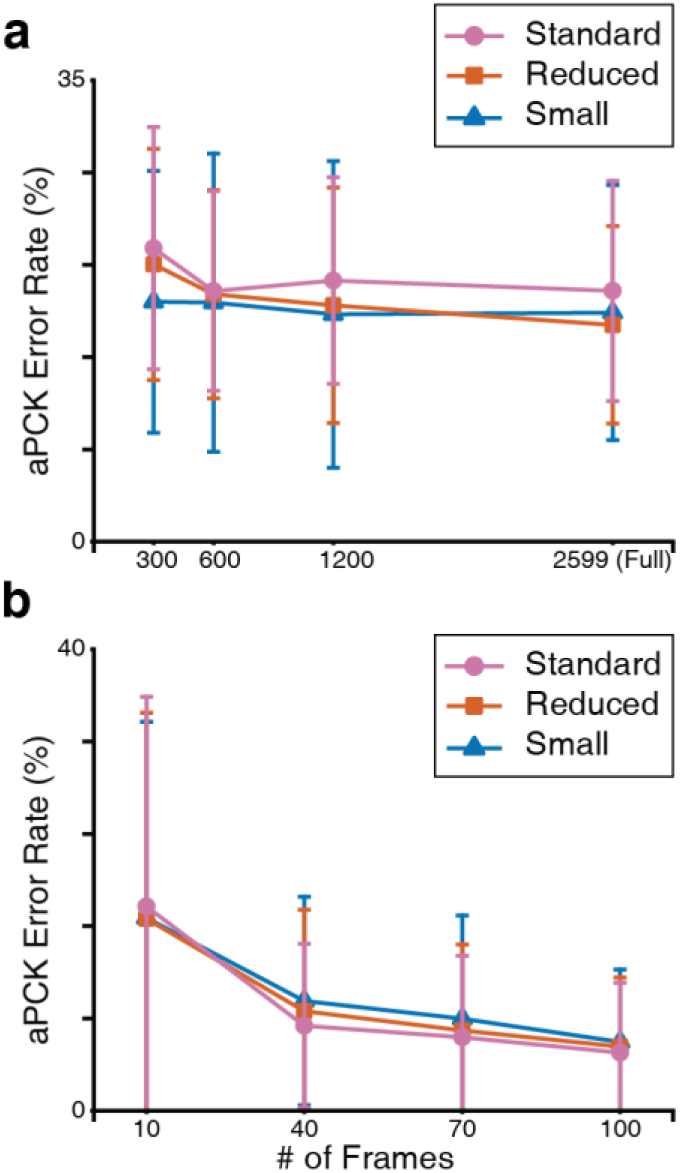
FlexibleBaseline model size evaluation. **a**, Performance on mouse side view dataset. **b**, Performance on fruit fly dataset. To simulate hardware constraints, all mouse side view models were trained for 40,000 (20×2,000) steps at a batch size of 10, and all fruit fly models were trained for 8,000 (10×800) steps at a batch size of 10.

### Improving Robustness Against Temporary Obstruction of OptiFlex with OpticalFlow

The OpticalFlow module morphs heatmap predictions of neighbouring frames onto the target frame using the Lucas-Kanade method^31^, implemented with the Farneback algorithm^32^ in OpenCV^33^. This morphed information is aggregated by a 3D convolution layer that essentially takes the weighted sum of all the morphed heatmaps (**Fig. 1b**). This implies that even if some key points are not visible in the target frame, the morphed information from nearby frames still provides sufficient information about the most likely location of the key points for the target frame. The morphed heatmaps from nearby frames can thus be considered as temporal context.

Of all the training datasets, temporary obstruction occurred most frequently in the mouse side view as the paws frequently overlap in this view. Therefore, contrasting mouse side view prediction results from before and after applying the OpticalFlow model best demonstrates robustness against temporary obstruction (**Fig. 5** and **Video S2**). We applied this procedure not only to FlexibleBaseline, but also to LEAP, DeepLabCut, and StackedDenseNet base models. Even though the OpticalFlow model integrated with FlexibleBaseline (i.e. OptiFlex) showed the best results, addition of the OpticalFlow model improved performance of all available base models (**Fig. 5a**). Results of OpticalFlow corrections are most evident through smoothing of trace curves of the keypoints. Sharp spikes in the trace curves were detected as model prediction errors, and the OpticalFlow curve comparisons in **Fig. 5b** and **Fig. 5c** provide good examples of where the spikes are smoothed out by OpticalFlow, indicating error correction. It should be noted that when the base model makes multiple consecutive erroneous predictions, the OpticalFlow model does not recognize those predictions as temporary obstructions, and does not make corrections (Hind Left Paw in **Fig. 5c**).

**Fig. 5.**
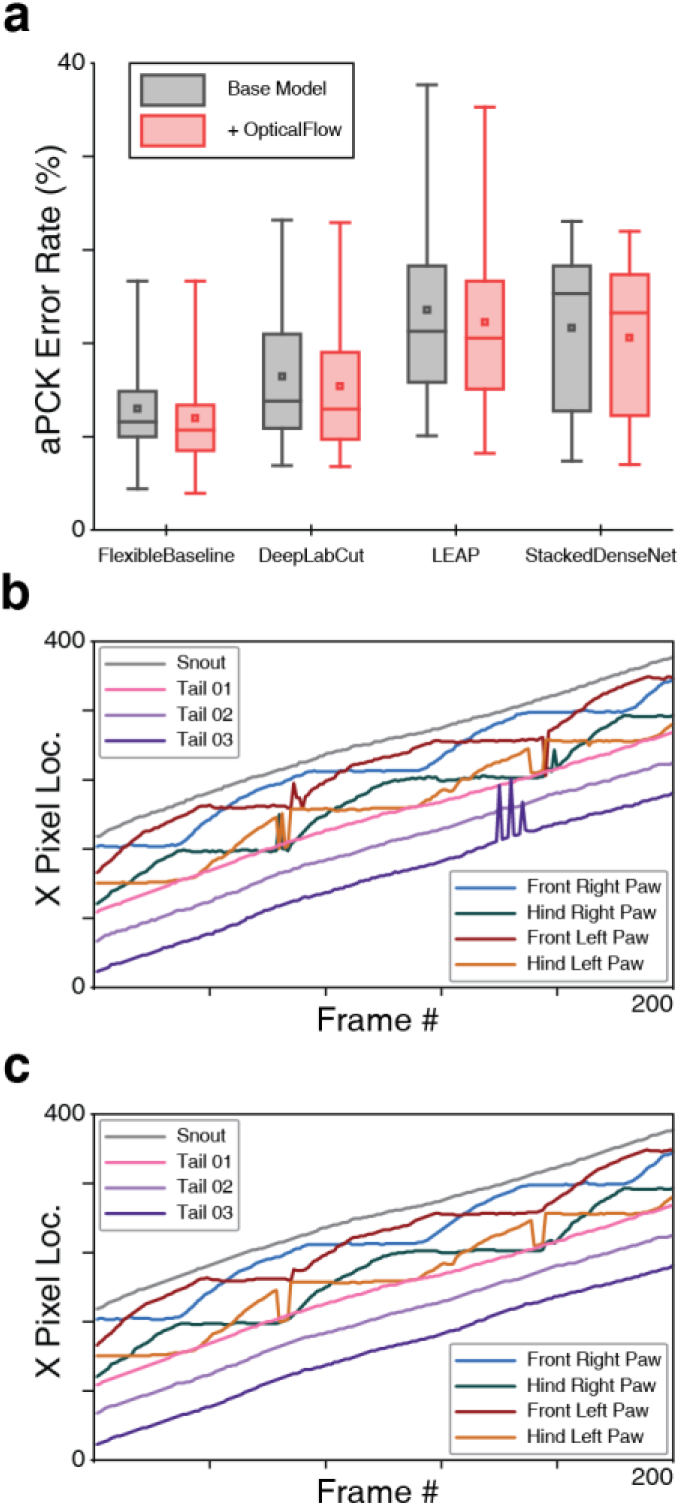
OpticalFlow model of OptiFlex evaluation. **a**, Box plot of test set prediction error rates of models with and without OpticalFlow grouped by base model. Box plot definition as specified in **Fig. 3a**. The benefits of the OpticalFlow model appeared universal in that it also improved the performance of the other base models (LEAP, DeepLabCut, and StackedDenseNet). **b**, X-value traces of FlexibleBaseline key point predictions for a mouse side view video (video code: 00000nst_0028) without OpticalFlow correction. **c**, X-value traces of OpticalFlow model key point predictions for the same mouse side view video (video code: 00000nst_0028, using FlexibleBaseline). Note that the differences after applying the OpticalFlow model were most prominently reflected in the smoothing of the trace curves of the key points. Sharp spikes in the trace curves corresponded to prediction errors detected by OpticalFlow.

### Exploration in Multi-View Enhancement

Instead of using multiple views to construct 3D representation of joint movements, we explored the idea of using heatmap predictions obtained from different views to correct each other. Exploiting multiple views of the same behaviour can improve the predictions, because certain features are more identifiable in one view than another and the geometrical configuration of the different views determines the information shared between them. To demonstrate the potential of multi-view correction, we developed a simple algorithm that corrects paw predictions in the mouse dataset using initial predictions from both the side and bottom views.

For the mouse dataset, the two perpendicular views (side and bottom) must share an axis, i.e. the x-axis, in 3D space. Based on analyses of the single view prediction results presented above, it could be determined that the bottom view model better predicted the position of the paws. As a consequence, the x-value of the paws from the bottom could be used as a reference to search for alternative prediction locations for the paws in the side view. These alternative prediction locations were generated by finding local maximums in the prediction heatmap using Gaussian filters in the side view. Finally, the optimised locations for the paws in the side view corresponded to the locations with the least difference in x-value from their respective key points in the reference (bottom) view.

The effects of the multi-view correction algorithm are demonstrated in **Fig. 6**, **Video S3** and **Table S8**. This algorithm leads to significant improvements on all base models, and a nearly perfect result is achieved after applying the algorithm to OptiFlex, which now integrates FlexibleBaseline, OpticalFlow, and multi-view corrections.

**Fig. 6.**
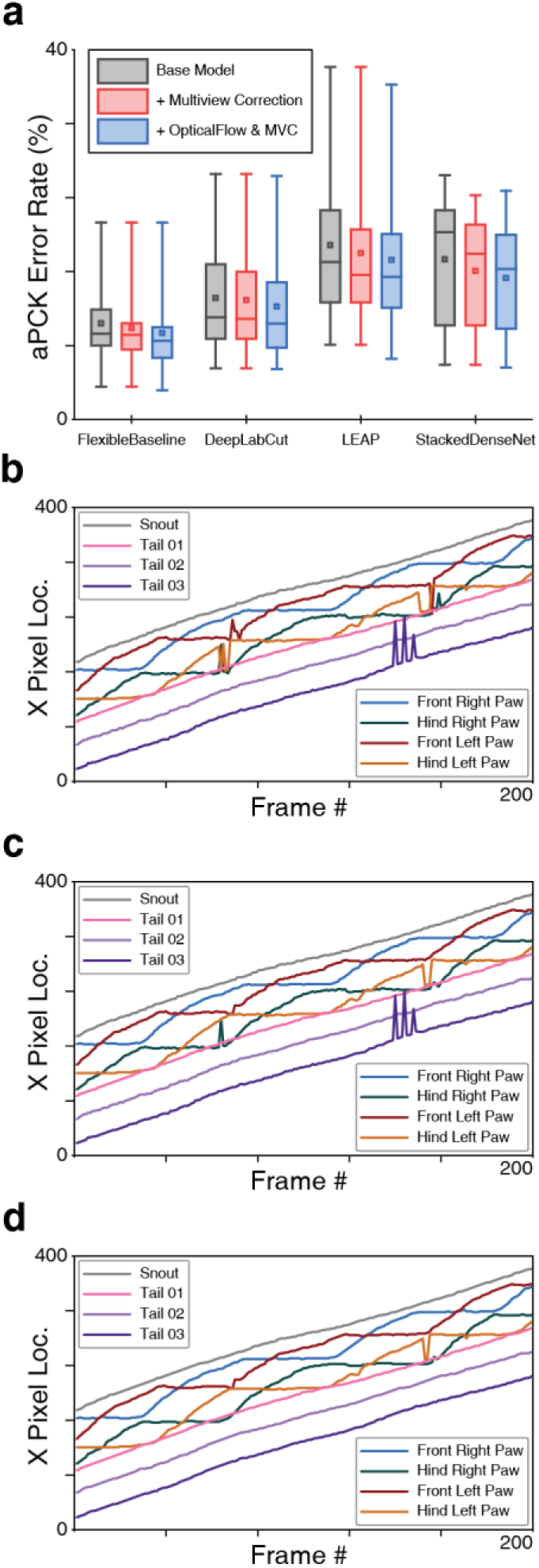
Multiview paw correction algorithm evaluation of OptiFlex. **a**, Box plot of test set prediction error rates of models with and without multi-view correction on paws, grouped by base model. Box plot definition as specified in **Fig. 3a**. **b**, X-value traces of FlexibleBaseline paw predictions for a mouse side view video (video code: 00000nst_0028_tst). **c**, X-value traces of FlexibleBaseline paw predictions for a mouse side view video (video code: 00000nst_0028_tst) after multi-view correction. **d**, X-value traces of OptiFlex, which integrates FlexibleBaseline and OpticalFlow, paw predictions for the same mouse side view video (video code: 00000nst_0028_tst) after multi-view correction.

## Discussion

This paper introduces the first video-based architecture for animal pose estimation, which we refer to as OptiFlex. We exploit a new intuitive metric, i.e. the percentage of correct key points or aPCK, to evaluate performance in the context of animal pose estimation. Based on analyses of behavioural experiments in four different animal species, OptiFlex achieved the lowest prediction error rates compared to other commonly used models for animal pose estimation. Our architecture leveraged the temporal context information through optical flow to enhance our FlexibleBaseline model, allowing this architecture to correct for temporarily obstructed key points. Moreover, the architecture also facilitated exploitation of multi-view analysis, further reducing prediction errors. Overall, the effectiveness and robustness of our OptiFlex architecture, which is available through our open source Github repository (https://github.com/saptera/OptiFlex), make it a potentially valuable component in any system that wishes to track animal behaviour.

### Dataset Considerations

With diverse body plans of different species and distinct experimental setups, each of the datasets used in our model assessments posed unique challenges. The mouse dataset was obtained on a setup with both side and bottom views, allowing the combination of spatial geometric information from both views. In the mouse side view dataset, it was challenging to continuously track mouse paw movements during locomotion as the paws are constantly alternating, often leading to temporary obstruction. This made the mouse dataset a perfect test case for handling temporary obstruction of key points.

The fruit fly dataset on the other hand, which was from the same dataset as used by LEAP^24^, had the largest number of key points to track, making models for this dataset more memory intensive than models of other datasets. At the same time, the tracking process was simplified by removal of image backgrounds and high visibility of the key points. The monkey dataset comprised only facial features, with one of the key points, the tongue, having a very low rate of occurrence. This feature made the monkey dataset a proper example for testing datasets with highly imbalanced key point occurrences. The zebrafish dataset, finally, was the only dataset that required tracking of multiple animals at the same time. This endeavour was particularly challenging as individual zebrafish are hard to distinguish, while they frequently traverse the field of view. Currently, OptiFlex can only track multiple animals under constraints such as a predetermined number of animals and consistent animal body shape throughout the dataset. This ability could be generalized to tracking an arbitrary number of animals by adding an extra object detection module, which is a potential direction for future work. Together, the data of the different species comprised a rich and diverse set of behavioural measurements that allowed us to test the limitations of OptiFlex and competing models to their full extent.

### Temporary Obstruction

Applying OpticalFlow turned out to be beneficial not only for FlexibleBaseline of OptiFlex, but also for all other base models tested here. In all cases, it was particularly instrumental in correcting for temporary obstruction. The issue of temporary obstruction has been identified before by others, but it has been partly circumvented by using applying different strategies^18, 24, 34^. For example, some models analysed selected datasets with a relatively high visibility of the key points^18, 34^, while others reported error distances at the 90^th^ percentile level even though the occurrence of obstructions was probably less than 10%^24^.

### Spatial, Geometrical, and Temporal Context

Fundamentally, animal key point movements happen in 3D space over time, making the intrinsic information four dimensional. If we attempt to make predictions on animal key point movements using only two dimensional data, such as a single video frame, then we have forfeited 2 dimensions of the information available. This fundamental flaw will persist despite improvements in models or training datasets. In the current paper, we demonstrated ways to take all 4 dimensions into account by using FlexibleBaseline to generate predictions based on 2D video frames, combining multi-view analysis to generate predictions in a 3D geometric context, and adding optical flow to utilize the fourth temporal dimension. By integrating all these features into a single architecture, OptiFlex provides the next step forward to use advanced deep learning tools to analyse animal behaviour non-invasively at a high spatiotemporal resolution.

## Methods

### Formulation

The goal of OptiFlex is to produce a set of heatmaps representing the likelihood of each key point appearing at each location of the image. We denote the pixel location of the *p*^*th*^ key point as *y*_*p*_ ∈ *Z* ⊂ ℝ^2^, where *Z* is the set of all (*x*, *y*) locations in an image and *p* ∈ {1 … *P*}.

The base model will have *N* outputs, each considered as a function. With the *i*^*th*^ output denoted by *b*_*i*_(⋅), where *i* ∈ {1 … *N*}. Usually, there are 1 to 2 outputs from the base model. Each output goes through a resizing process (usually deconvolution), denoted *d*_*i*_(⋅), to produce a set of intermediate heatmaps ***h***_*i*_. Thus, if we let the input frame be denoted as **x**, we have 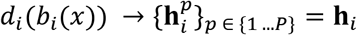, where 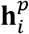 is the heatmap for the *p*^*th*^ key point from the *i*^*th*^ output. The intermediate heatmaps ***h***_*i*_ is used to compare against labels for intermediate supervision, and the final set of heatmaps ***h***_*N*_, denoted **f**, is used as the output heatmap for the base model.

To use context information from surrounding frames, the base model is first applied to predict heatmaps for all surrounding frames. Let {**x**^k^}_k_ ∈ {*t*−*s*×*f* …*t* …*t*+*s*×*f*}be a sequence of input frames, where *t* is the index of the target frame, *s* is the skip ratio, and **f** is the frame range. We apply the base model to each frame to get the output heatmaps *d*_*N*_(*g*_*N*_(**x**^k^)) = **f**^k^. Next, we compute optical flow between each of the output heatmaps of surrounding frames and target frame **x**^*t*^. We denote the optical flow function as ϕ(⋅,⋅), which outputs optical flow morphed heatmaps of input heatmaps **f**^k^ with reference to target frame heatmaps **f**^*t*^: *ϕ*(**f**^k^, **f**^*t*^) = **o**^k^. Note that **f**^*t*^ = *ϕ*(**f**^*t*^, **f**^*t*^) = **o**^*t*^. We pass all of the optical flow morphed heatmaps through a 1×1 convolution layer to get the final output heatmaps of the entire model: **y** = *conv*_1*x*1_ ({**o**^k^}k ∈ {*t*−*s*×*f* …*t* …*t*+*s*×*f*}). This 1×1 convolution essentially acts as a weighted sum of all the optical flow morphed heatmaps. Finally, we can get the predicted pixel location of the *p*^*th*^ key point, denoted 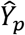, by getting the global maximum of the *p*^*th*^ final output heatmap **y**_*p*_. aPCK calculations can be done by comparing 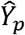 against *y*_*p*_.

### Datasets

Our multi-species datasets cover some commonly used animal body plans in locomotion experiments: mouse (*Mus musculus*), zebrafish (*Danio rerio*), fruit fly (*Drosophila melanogaster*), and monkey (*Rhesus macaque*). The mouse dataset was acquired on a LocoMouse setup^35^, which contains a straight corridor and a bottom mirror that permits observation of side and bottom views of the mouse with a single camera. The zebrafish dataset was recorded with a camera mounted above a fish tank to film the activity of multiple fish. The fruit fly dataset was downloaded from the Princeton Neuroscience Institute (http://arks.princeton.edu/ark:/88435/dsp01pz50gz79z)^24^. The monkey dataset was obtained with a camera filming the facial behaviour of a rhesus monkey, with an installed lick port.

### Dataset Preprocessing

The raw dataset is first split into train, validation, and test sets, with roughly a 3꞉1꞉1 ratio. After the split, the validation and test sets are ready after resizing to the target dimension, while the train set requires further processing. Three major steps were performed on the train set of each dataset before being used for the data generators of the base models: augment, resize image and labels, and label conversion.

1. Augment all images and labels within the training set with random rotation and flipping. Images are kept without cropping and padded with black background, thus retaining all image information. For our datasets, the angle range for random rotation is (−10°, 10°). No flipping is applied to the mouse or fruit fly train sets; the monkey train set is randomly flipped about the y-axis, and zebrafish train set is randomly flipped about all axes (x-axis, y-axis, xy-axis).
2. Resize all augmented images and labels to the same size. Images were resampled using pixel area relation, a preferred method for decimating images^33^. The same transformations were also applied to the labels.
3. Convert human defined labels to heatmaps. Human defined labels are either pixel coordinates or bounding boxes. For the heatmap of each key point, a 2D tensor of image size is initialized with all zeros. The heatmap is image size, so small key points such as mouse paws do not get shrunk to a single pixel. A 2D Gaussian distribution is generated by probability density function (PDF) within a user defined area or the bounding box area on the tensor, with the center of the area being the ground truth location for the joint and location of peak value for the 2D Gaussian distribution. Then, the heatmap is normalized to make the maximum value 1.0, and any value on the heatmap smaller than 0.1 is set to 0. The third step is the most crucial step as it is closely tied to the eventual evaluation using aPCK, because the area defined by the labeller for the 2D Gaussian distribution will be the area considered accepted or “correct”.

Since the datasets were too large to be directly stored in memory, train and validation sets were converted to data generators before feeding them into the model. For each base model, a single training input consists of a batch of images converted to tensors and a batch of multiple copies of label heatmap tensor, depending on number of stages. The tensors are multiplied by a user defined peak value to increase contrast between label region and remaining pixels. From our training experience, most models cannot be trained without this process.

To train the OpticalFlow model, we first need an ordered sequence of 2*n* + 1 images to be used as inputs to a pre-trained base model. The base model produces an ordered sequence of 2*n* + 1 heatmap tensors as output, which is then fed into the OpticalFlow component and the labels for the OpticalFlow training process will be a single set of heatmap labels.

### Base Model Training Setup

To make fair comparisons between various base model designs, we set the training length at 50 epochs and the batch size at 10 for all base models on all datasets. We also tried to keep a constant learning rate of 0.0001 across different models and datasets. However, when using the LEAP model, a learning rate of 0.0001 led to wrong predictions on one of the key points for the mouse bottom view dataset. We therefore used a learning rate of 0.0003 instead in this specific instance. All heatmap labels had a truncated normal distribution with peak value of 16 located at the manually labeled key point position. All models were trained using an ADAM^36^ optimizer with beta1=0.9, beta2=0.999, and no decay.

### Base Model Implementation

Again, to make fair comparisons between base model designs, all training, data generation and evaluation procedures were identical. All base models were implemented using Keras^37^ and are available on Github.

The standard FlexibleBaseline models used in model comparisons all had the same hyperparameter: ImageNet pre-trained ResNet50 backbone outputs after Conv4 block and the filter number for the last 3 transposed convolution layers are 64, 64, and 2× number of key points respectively. There is an intermediate supervision after Conv3 block of ResNet50 backbone.

LEAP models were implemented exactly based on the specification of the original paper. Since LEAP also produces heatmaps of original image size, our data generation process worked perfectly with the model.

DeepLabCut models were also implemented according to the original paper, except the original hyperparameters produced prediction heatmaps were smaller than the original images. For DeepLabCut to train using the same data generation process, we changed the kernel size and stride of the final transpose convolution layer to 36×36 and 32×32 respectively.

DeepPoseKit was originally implemented using Keras^18^, so our StackedDenseNet implementation was nearly identical to the DeepPoseKit Github implementation, with some minor refactoring to ensure the model works with the rest of the code base. Since the original Stacked DenseNet also produced prediction heatmaps of a smaller size, an additional TransitionUp module (from DenseNet) was added before each output layer to ensure the model produce output of original image size.

### Flexibility Comparison Setup

The 3 versions of FlexibleBaseline had hyperparameters specified in **Table S3**. For fruit fly datasets, all model and dataset combinations were trained with 8,000 images randomly sampled with replacement from their respective training set. For mouse side view datasets, all model and dataset combinations are trained with 400,000 images randomly sampled with replacement from their respective training set.

### OpticalFlow Implementation and Setup

Our OpticalFlow model was similar in principle to a component of Flowing ConvNet^38^, but had significant changes in implementation to allow for skip ratio and predefined frame range. The hyperparameter values for OpticalFlow from Farneback algorithm are: window size of 27 pixels, pyramid scale of 0.5 with 5 levels and 8 iteration on each pyramid level; pixel neighbourhood size was 7 for polynomial expansion, with a corresponding *poly_sigma* of 1.5.

In our OpticalFlow models comparisons, all OpticalFlow models were trained for 30 epochs with a skip ratio of 1. All OpticalFlow models had a learning rate of 0.0001, except for StackedDenseNet, which had a learning rate of 0.00015. The OpticalFlow model for StackedDenseNet used a slightly higher learning rate because its validation curve did not plateau with learning rate 0.0001 after 30 epochs. All OpticalFlow models had a frame range of 4, except for LEAP, which had a frame range of 2. The LEAP base model had many more prediction errors than other base models, so including more frames often introduced false information to the target frame.

### Computing Environment

All training and inference were done on VM instances from Computing Engine of Google Cloud with identical configuration. Each VM instance was a general-purpose N1 series machine with 24 vCPU,156GB of memory and 2 Nvidia Tesla V100 16GB VRAM GPU. The OS image on each instance was “Deep Learning Image: TensorFlow 1.13.1 m27” with CUDA 10.0 installed.

## Supporting information

Vid-S1a

Vid-S1b

Vid-S1c

Vid-S1d

Vid-S2

Vid-S3

## Notes

#### Summary of Updates

Added URL for our Github Repository

https://github.com/saptera/OptiFlex

