## Supplemental Materials for "OptiFlex: video-based animal pose estimation using deep learning enhanced by optical flow"

**Table S1, Dataset attributes**

**Table S2, Training dataset division**

**Table S3, Hyperparameters of FlexibleBaseline versions**

**Table S4, Hyperparameters for Optical flow models**

**Table S5-a~e, Model prediction aPCK accuracy (%) on different datasets.** Best results at each joint is marked as GREEN, best statistical results is marked as **GREEN_BOLD**.

**Table S6-a~b, Comparison on FlexibleBaseline model size prediction aPCK accuracy (%) on [mouse side-view] and [fruit fly] datasets.** Best results are calculated separately by the group of data size. Best results at each joint is marked as GREEN, best statistical results is marked as **GREEN_BOLD**.

**Table S7. Comparison between BaseModel and OpticalFlow prediction aPCK accuracy (%) on [mouse side-view] dataset.** Best results at each joint is marked as GREEN, best statistical results is marked as **GREEN_BOLD**.

**Table S8. Comparison between BaseModel and MultiviewCorrection prediction aPCK accuracy (%) on [mouse side-view] dataset.** Best results at each joint is marked as GREEN, best statistical results is marked as **GREEN_BOLD**.

**Table S9 and Fig. S1, Comparison on fully trained FlexibleBaseline model size prediction evaluation on [mouse side-view] dataset.** Best results at each joint is marked as GREEN, best statistical results is marked as **GREEN_BOLD**.

**Fig. S2, Manual labelling GUI**

**Fig. S3, Dataset creation GUI**

**Fig. S4, Dataset example**

**Video S1-a~d, Compare prediction results of different base models**

**Video S2, Compare prediction results of FlexibleBaseline /w OpticalFlow**

**Video S3, Compare prediction results of FlexibleBaseline /w MultiviewCorrection**

### Table S1 | Dataset attributes

| Dataset | Image Size (w × h) | Frame Rate (fps) | Total # of Frames | # of Key Points |
|---|---|---|---|---|
| Mouse (Side) | 512 × 128 | 400 reduced to 100 | 5134 | 8 |
| Mouse (Bottom) | 512 × 128 | 400 reduced to 100 | 5134 | 8 |
| Fruit Fly | 512 × 256 | 100 | 1500 | 32 |
| Monkey | 512 × 256 | 40 | 391 | 7 |
| Zebrafish | 512 × 256 | 25 | 451 | 12 |

### Table S2 | Training dataset division

| Dataset | Train Set Image # | Validation Set Image # | Test Set Image # |
|---|---|---|---|
| Mouse (Side) | 2599 | 1207 | 1051 |
| Mouse (Bottom) | 2606 | 1325 | 1203 |
| Fruit Fly | 900 | 300 | 300 |
| Monkey | 235 | 78 | 78 |
| Zebrafish | 271 | 90 | 90 |

*All images in train set were augmented 8 times. The table shows the number of images before augmentation.*

### Table S3 | Hyperparameters of FlexibleBaseline versions

| FlexibleBaseline Version | ResNet50 Backbone Output | Intermediate Supervision | # Filters in 1st TrsConv | # Filters in 2nd TrsConv | # Filters in 3rd TrsConv |
|---|---|---|---|---|---|
| Standard | after Conv4 | after Conv3 | 64 | 64 | 2x # key points |
| Reduced | after Conv3 | after Conv2 | 64 | 64 | 2x # key points |
| Small | after Conv2 | N/A | 32 | 32 | 2x # key points |

*Conv stands for "Convolutional Layer"; TrsConv stands for "Transposed Convolutional Layer".*

### Table S4 | Hyperparameters for Optical flow models

| Hyperparameter | FlexibleBaseline | DeepLabCut | LEAP | StackedDenseNet |
|---|---|---|---|---|
| Skip Ratio | 1 | 1 | 1 | 1 |
| Frame Range | 4 | 4 | 2 | 4 |
| Learning Rate | 0.0001 | 0.0001 | 0.0001 | 0.00015 |

### Table S5-a | Model prediction aPCK accuracy (%) on [mouse side-view] dataset

| Joint Name | FlexibleBaseline | DeepLabCut | LEAP | StackedDenseNet |
|---|---|---|---|---|
| Front Right Paw | 0.923882 | 0.913416 | 0.785918 | 0.735490 |
| Hind Right Paw | 0.888677 | 0.868696 | 0.830637 | 0.789724 |
| Front Left Paw | 0.873454 | 0.836346 | 0.761180 | 0.801142 |
| Hind Left Paw | 0.907707 | 0.827783 | 0.828735 | 0.793530 |
| Snout | 0.906755 | 0.910561 | 0.874405 | 0.896289 |
| Tail 01 | 0.964795 | 0.944814 | 0.872502 | 0.941009 |
| Tail 02 | 0.786870 | 0.734539 | 0.618459 | 0.757374 |
| Tail 03 | 0.916270 | 0.912464 | 0.919125 | 0.900095 |
| ***COUNT*** | **6** | 1 | 1 | 0 |
| ***MEAN*** | **0.896051** | 0.868577 | 0.811370 | 0.826832 |
| ***SD*** | **0.051610** | 0.067803 | 0.092872 | 0.075159 |

### Table S5-b | Model prediction aPCK accuracy (%) on [mouse bottom-view] dataset

| Joint Name | FlexibleBaseline | DeepLabCut | LEAP | StackedDenseNet |
|---|---|---|---|---|
| Front Right Paw | 0.932668 | 0.915212 | 0.914381 | 0.935162 |
| Hind Right Paw | 0.981712 | 0.983375 | 0.970906 | 0.980881 |
| Front Left Paw | 0.968412 | 0.951787 | 0.912718 | 0.944306 |
| Hind Left Paw | 0.975894 | 0.955943 | 0.940150 | 0.935162 |
| Snout | 0.733998 | 0.721530 | 0.713217 | 0.674148 |
| Tail 01 | 0.975894 | 0.971737 | 0.949293 | 0.914381 |
| Tail 02 | 0.714048 | 0.710723 | 0.551122 | 0.754780 |
| Tail 03 | 0.827099 | 0.814630 | 0.768911 | 0.847049 |
| ***COUNT*** | **4** | 1 | 0 | 3 |
| ***MEAN*** | **0.888716** | 0.878117 | 0.840087 | 0.873234 |
| ***SD*** | 0.113625 | 0.112932 | 0.148569 | **0.107141** |

**Table S5-c | Model prediction aPCK accuracy (%) on [fruit fly] dataset**

| Joint Name | FlexibleBaseline | DeepLabCut | LEAP | StackedDenseNet |
|---|---|---|---|---|
| head | 1.000000 | 0.996667 | 1.000000 | 1.000000 |
| eyeL | 1.000000 | 0.996667 | 1.000000 | 1.000000 |
| eyeR | 1.000000 | 0.996667 | 1.000000 | 1.000000 |
| neck | 1.000000 | 0.996667 | 1.000000 | 0.996667 |
| thorax | 1.000000 | 0.996667 | 1.000000 | 0.996667 |
| abdomen | 1.000000 | 1.000000 | 1.000000 | 0.996667 |
| forelegR1 | 0.993333 | 0.996667 | 0.996667 | 0.993333 |
| forelegR2 | 0.976667 | 0.970000 | 0.966667 | 0.976667 |
| forelegR3 | 0.950000 | 0.956667 | 0.943333 | 0.953333 |
| forelegR4 | 0.956667 | 0.956667 | 0.953333 | 0.953333 |
| midlegR1 | 0.986667 | 0.986667 | 0.983333 | 0.983333 |
| midlegR2 | 0.993333 | 0.986667 | 0.993333 | 0.983333 |
| midlegR3 | 0.986667 | 0.986667 | 0.976667 | 0.986667 |
| midlegR4 | 0.976667 | 0.976667 | 0.943333 | 0.966667 |
| hindlegR1 | 0.993333 | 0.996667 | 0.993333 | 0.996667 |
| hindlegR2 | 0.943333 | 0.940000 | 0.933333 | 0.943333 |
| hindlegR3 | 0.933333 | 0.913333 | 0.910000 | 0.916667 |
| hindlegR4 | 0.860000 | 0.860000 | 0.810000 | 0.810000 |
| forelegL1 | 0.996667 | 0.996667 | 0.996667 | 0.993333 |
| forelegL2 | 0.966667 | 0.966667 | 0.960000 | 0.966667 |
| forelegL3 | 0.963333 | 0.963333 | 0.953333 | 0.970000 |
| forelegL4 | 0.963333 | 0.960000 | 0.936667 | 0.966667 |
| midlegL1 | 0.990000 | 0.993333 | 0.990000 | 0.993333 |
| midlegL2 | 0.976667 | 0.970000 | 0.963333 | 0.970000 |
| midlegL3 | 0.953333 | 0.966667 | 0.966667 | 0.960000 |
| midlegL4 | 0.966667 | 0.963333 | 0.946667 | 0.966667 |
| hindlegL1 | 0.990000 | 0.990000 | 0.990000 | 0.986667 |
| hindlegL2 | 0.940000 | 0.943333 | 0.943333 | 0.923333 |
| hindlegL3 | 0.950000 | 0.943333 | 0.933333 | 0.960000 |
| hindlegL4 | 0.910000 | 0.936667 | 0.866667 | 0.890000 |
| wingL | 1.000000 | 0.996667 | 0.993333 | 1.000000 |
| wingR | 1.000000 | 1.000000 | 0.993333 | 1.000000 |
| ***COUNT*** | **22** | 17 | 12 | 15 |
| ***MEAN*** | **0.972396** | 0.971875 | 0.963646 | 0.968750 |
| ***SD*** | 0.031557 | **0.030702** | 0.042429 | 0.039520 |

### Table S5-d | Model prediction aPCK accuracy (%) on [monkey] dataset

| Joint Name | FlexibleBaseline | DeepLabCut | LEAP | StackedDenseNet |
|---|---|---|---|---|
| upperlip1 | 1.000000 | 1.000000 | 1.000000 | 1.000000 |
| upperlip2 | 1.000000 | 1.000000 | 1.000000 | 1.000000 |
| lowerlip1 | 1.000000 | 1.000000 | 1.000000 | 1.000000 |
| lowerlip2 | 0.987179 | 0.987179 | 0.974359 | 0.987179 |
| brow | 1.000000 | 1.000000 | 1.000000 | 1.000000 |
| lickspout | 1.000000 | 1.000000 | 1.000000 | 1.000000 |
| tongue | 0.987179 | 0.987179 | 0.987179 | 0.961538 |
| ***COUNT*** | **7** | **7** | 6 | 6 |
| ***MEAN*** | **0.996337** | **0.996337** | 0.994505 | 0.992674 |
| ***SD*** | **0.006256** | **0.006256** | 0.010087 | 0.014537 |

### Table S5-e | Model prediction aPCK accuracy (%) on [zebrafish] dataset

| Joint Name | FlexibleBaseline | DeepLabCut | LEAP | StackedDenseNet |
|---|---|---|---|---|
| zf_01 | 0.888889 | 0.888889 | 0.866667 | 0.822222 |
| zf_02 | 0.922222 | 0.844444 | 0.666667 | 0.266667 |
| zf_03 | 0.900000 | 0.888889 | 0.411111 | 0.222222 |
| zf_04 | 0.755556 | 0.777778 | 0.388889 | 0.266667 |
| zf_05 | 0.966667 | 0.966667 | 0.844444 | 0.944444 |
| zf_06 | 0.811111 | 0.788889 | 0.566667 | 0.155556 |
| zf_07 | 0.777778 | 0.722222 | 0.377778 | 0.122222 |
| zf_08 | 0.911111 | 0.833333 | 0.622222 | 0.300000 |
| zf_09 | 0.944444 | 0.955556 | 0.711111 | 0.700000 |
| zf_10 | 0.844444 | 0.844444 | 0.655556 | 0.800000 |
| zf_11 | 0.944444 | 0.944444 | 0.655556 | 0.833333 |
| zf_12 | 0.877778 | 0.700000 | 0.388889 | 0.466667 |
| ***COUNT*** | **10** | 6 | 0 | 0 |
| ***MEAN*** | **0.878704** | 0.846296 | 0.596296 | 0.491667 |
| ***SD*** | **0.068076** | 0.087724 | 0.173065 | 0.306023 |

**Table S6-a | Comparison on FlexibleBaseline model size prediction aPCK accuracy (%) on [mouse side-view] dataset**

| Data Size | 300 | | | 600 | | | 1200 | | | 2599 (Full) | | |
|---|---|---|---|---|---|---|---|---|---|---|---|---|
| **Joint Name** | **Standard** | **Reduced** | **Small** | **Standard** | **Reduced** | **Small** | **Standard** | **Reduced** | **Small** | **Standard** | **Reduced** | **Small** |
| Front Right Paw | 0.712655 | 0.717412 | 0.771646 | 0.781161 | 0.739296 | 0.731684 | 0.800190 | 0.778306 | 0.750714 | 0.796384 | 0.784015 | 0.764034 |
| Hind Right Paw | 0.717412 | 0.719315 | 0.746908 | 0.786870 | 0.795433 | 0.788773 | 0.774500 | 0.770695 | 0.851570 | 0.809705 | 0.808754 | 0.815414 |
| Front Left Paw | 0.700285 | 0.712655 | 0.787821 | 0.779258 | 0.770695 | 0.822074 | 0.763083 | 0.784967 | 0.807802 | 0.784967 | 0.833492 | 0.811608 |
| Hind Left Paw | 0.741199 | 0.796384 | 0.823026 | 0.798287 | 0.783064 | 0.793530 | 0.777355 | 0.810657 | 0.833492 | 0.739296 | 0.816365 | 0.842055 |
| Snout | 0.891532 | 0.876308 | 0.922931 | 0.853473 | 0.886775 | 0.919125 | 0.907707 | 0.898192 | 0.900095 | 0.877260 | 0.882017 | 0.891532 |
| Tail 01 | 0.932445 | 0.937203 | 0.963844 | 0.960990 | 0.962892 | 0.969553 | 0.924833 | 0.972407 | 0.977165 | 0.957184 | 0.961941 | 0.965747 |
| Tail 02 | 0.704091 | 0.714558 | 0.656518 | 0.696480 | 0.728830 | 0.618459 | 0.686013 | 0.683159 | 0.594672 | 0.682207 | 0.712655 | 0.640343 |
| Tail 03 | 0.819220 | 0.843007 | 0.870599 | 0.822074 | 0.830637 | 0.905804 | 0.781161 | 0.867745 | 0.904853 | 0.830637 | 0.884872 | 0.879163 |
| ***COUNT*** | 0 | 1 | **7** | 2 | 2 | **4** | 3 | 0 | **5** | 1 | 3 | **4** |
| ***MEAN*** | 0.777355 | 0.789605 | **0.817912** | 0.809824 | 0.812203 | **0.818625** | 0.801855 | 0.820766 | **0.827545** | 0.809705 | **0.835514** | 0.826237 |
| ***SD*** | 0.091991 | **0.087755** | 0.099472 | **0.075753** | 0.079037 | 0.113161 | **0.078404** | 0.089304 | 0.116350 | 0.083456 | **0.075039** | 0.096758 |

### Table S6-b | Comparison on FlexibleBaseline model size prediction aPCK accuracy (%) on [fruit fly] dataset

| Data Size | 10 | | | 40 | | | 70 | | | 100 | | |
|---|---|---|---|---|---|---|---|---|---|---|---|---|
| **Joint Name** | **Standard** | **Reduced** | **Small** | **Standard** | **Reduced** | **Small** | **Standard** | **Reduced** | **Small** | **Standard** | **Reduced** | **Small** |
| head | 1.000000 | 1.000000 | 1.000000 | 1.000000 | 1.000000 | 1.000000 | 1.000000 | 1.000000 | 1.000000 | 1.000000 | 1.000000 | 1.000000 |
| eyeL | 1.000000 | 1.000000 | 1.000000 | 1.000000 | 1.000000 | 1.000000 | 1.000000 | 1.000000 | 1.000000 | 1.000000 | 1.000000 | 1.000000 |
| eyeR | 1.000000 | 1.000000 | 1.000000 | 1.000000 | 1.000000 | 0.983333 | 1.000000 | 1.000000 | 1.000000 | 1.000000 | 1.000000 | 1.000000 |
| neck | 1.000000 | 1.000000 | 1.000000 | 1.000000 | 1.000000 | 1.000000 | 1.000000 | 1.000000 | 1.000000 | 1.000000 | 1.000000 | 1.000000 |
| thorax | 1.000000 | 1.000000 | 0.996667 | 1.000000 | 1.000000 | 0.996667 | 1.000000 | 1.000000 | 0.996667 | 1.000000 | 1.000000 | 1.000000 |
| abdomen | 0.993333 | 1.000000 | 1.000000 | 1.000000 | 1.000000 | 1.000000 | 1.000000 | 1.000000 | 1.000000 | 1.000000 | 1.000000 | 1.000000 |
| forelegR1 | 0.993333 | 0.990000 | 0.990000 | 0.993333 | 0.993333 | 0.993333 | 0.990000 | 0.990000 | 0.993333 | 0.993333 | 0.993333 | 0.990000 |
| forelegR2 | 0.880000 | 0.870000 | 0.796667 | 0.890000 | 0.876667 | 0.876667 | 0.916667 | 0.906667 | 0.860000 | 0.940000 | 0.946667 | 0.933333 |
| forelegR3 | 0.703333 | 0.686667 | 0.640000 | 0.840000 | 0.796667 | 0.746667 | 0.880000 | 0.836667 | 0.836667 | 0.923333 | 0.900000 | 0.906667 |
| forelegR4 | 0.630000 | 0.743333 | 0.766667 | 0.930000 | 0.890000 | 0.903333 | 0.930000 | 0.893333 | 0.916667 | 0.943333 | 0.920000 | 0.916667 |
| midlegR1 | 0.970000 | 0.956667 | 0.960000 | 0.983333 | 0.976667 | 0.980000 | 0.983333 | 0.970000 | 0.980000 | 0.980000 | 0.980000 | 0.973333 |
| midlegR2 | 0.813333 | 0.916667 | 0.863333 | 0.933333 | 0.980000 | 0.946667 | 0.963333 | 0.970000 | 0.980000 | 0.963333 | 0.963333 | 0.963333 |
| midlegR3 | 0.636667 | 0.630000 | 0.556667 | 0.846667 | 0.903333 | 0.870000 | 0.913333 | 0.946667 | 0.966667 | 0.960000 | 0.956667 | 0.963333 |
| midlegR4 | 0.613333 | 0.756667 | 0.723333 | 0.900000 | 0.890000 | 0.876667 | 0.923333 | 0.893333 | 0.916667 | 0.930000 | 0.936667 | 0.920000 |
| hindlegR1 | 0.990000 | 0.990000 | 0.976667 | 0.990000 | 0.983333 | 0.986667 | 0.993333 | 0.996667 | 0.986667 | 0.993333 | 0.996667 | 0.980000 |
| hindlegR2 | 0.740000 | 0.660000 | 0.620000 | 0.876667 | 0.886667 | 0.866667 | 0.890000 | 0.876667 | 0.910000 | 0.930000 | 0.916667 | 0.903333 |
| hindlegR3 | 0.570000 | 0.723333 | 0.683333 | 0.853333 | 0.730000 | 0.810000 | 0.873333 | 0.843333 | 0.760000 | 0.886667 | 0.813333 | 0.843333 |
| hindlegR4 | 0.386667 | 0.323333 | 0.366667 | 0.736667 | 0.706667 | 0.713333 | 0.686667 | 0.693333 | 0.633333 | 0.700000 | 0.736667 | 0.720000 |
| forelegL1 | 0.996667 | 0.993333 | 0.990000 | 0.996667 | 0.996667 | 0.996667 | 0.996667 | 0.996667 | 0.993333 | 0.996667 | 0.996667 | 0.990000 |
| forelegL2 | 0.850000 | 0.866667 | 0.793333 | 0.873333 | 0.913333 | 0.843333 | 0.863333 | 0.936667 | 0.930000 | 0.950000 | 0.936667 | 0.920000 |
| forelegL3 | 0.766667 | 0.753333 | 0.766667 | 0.880000 | 0.833333 | 0.843333 | 0.913333 | 0.883333 | 0.856667 | 0.923333 | 0.933333 | 0.903333 |
| forelegL4 | 0.746667 | 0.846667 | 0.786667 | 0.936667 | 0.883333 | 0.860000 | 0.920000 | 0.936667 | 0.886667 | 0.940000 | 0.943333 | 0.946667 |
| midlegL1 | 0.983333 | 0.980000 | 0.980000 | 0.986667 | 0.990000 | 0.986667 | 0.993333 | 0.990000 | 0.983333 | 0.990000 | 0.990000 | 0.990000 |
| midlegL2 | 0.850000 | 0.920000 | 0.880000 | 0.946667 | 0.930000 | 0.916667 | 0.956667 | 0.953333 | 0.900000 | 0.956667 | 0.943333 | 0.946667 |
| midlegL3 | 0.593333 | 0.586667 | 0.580000 | 0.890000 | 0.873333 | 0.883333 | 0.910000 | 0.910000 | 0.906667 | 0.933333 | 0.930000 | 0.913333 |
| midlegL4 | 0.830000 | 0.780000 | 0.820000 | 0.880000 | 0.833333 | 0.873333 | 0.923333 | 0.890000 | 0.893333 | 0.930000 | 0.933333 | 0.916667 |
| hindlegL1 | 0.990000 | 0.990000 | 0.990000 | 0.990000 | 0.976667 | 0.986667 | 0.990000 | 0.990000 | 0.970000 | 0.990000 | 0.990000 | 0.986667 |
| hindlegL2 | 0.790000 | 0.670000 | 0.806667 | 0.866667 | 0.800000 | 0.783333 | 0.900000 | 0.873333 | 0.863333 | 0.913333 | 0.906667 | 0.880000 |
| hindlegL3 | 0.523333 | 0.553333 | 0.773333 | 0.840000 | 0.830000 | 0.786667 | 0.876667 | 0.826667 | 0.780000 | 0.910000 | 0.876667 | 0.883333 |
| hindlegL4 | 0.540000 | 0.553333 | 0.586667 | 0.813333 | 0.783333 | 0.716667 | 0.793333 | 0.803333 | 0.773333 | 0.836667 | 0.853333 | 0.830000 |
| wingL | 0.980000 | 0.973333 | 0.986667 | 0.980000 | 0.990000 | 0.946667 | 0.990000 | 0.983333 | 0.986667 | 0.980000 | 0.956667 | 0.986667 |
| wingR | 0.976667 | 0.963333 | 0.973333 | 0.993333 | 0.990000 | 0.983333 | 0.996667 | 0.986667 | 0.993333 | 0.996667 | 0.976667 | 0.986667 |
| ***COUNT*** | **20** | 14 | 12 | **26** | 14 | 6 | **23** | 14 | 9 | **22** | 19 | 11 |
| ***MEAN*** | 0.823021 | **0.833646** | 0.832917 | **0.926458** | 0.913646 | 0.904896 | **0.936458** | 0.930521 | 0.920417 | **0.949687** | 0.944583 | 0.940417 |
| ***SD*** | 0.181863 | 0.178601 | **0.170276** | **0.071105** | 0.087778 | 0.090292 | **0.070746** | 0.074416 | 0.089454 | 0.060608 | **0.060130** | 0.062892 |

### Table S7 | Comparison between BaseModel and OpticalFlow prediction aPCK accuracy (%) on [mouse side-view] dataset

| Joint Name | FlexibleBaseline | | DeepLabCut | | LEAP | | StackedDenseNet | |
|---|---|---|---|---|---|---|---|---|
| | BaseModel | OpticalFlow | BaseModel | OpticalFlow | BaseModel | OpticalFlow | BaseModel | OpticalFlow |
| Front Right Paw | 0.923882 | 0.929591 | 0.913416 | 0.914367 | 0.785918 | 0.793530 | 0.735490 | 0.744053 |
| Hind Right Paw | 0.888677 | 0.900095 | 0.868696 | 0.878211 | 0.830637 | 0.832540 | 0.789724 | 0.809705 |
| Front Left Paw | 0.873454 | 0.885823 | 0.836346 | 0.848716 | 0.761180 | 0.780209 | 0.801142 | 0.814462 |
| Hind Left Paw | 0.907707 | 0.914367 | 0.827783 | 0.846813 | 0.828735 | 0.838249 | 0.793530 | 0.813511 |
| Snout | 0.906755 | 0.914367 | 0.910561 | 0.915319 | 0.874405 | 0.887726 | 0.896289 | 0.902950 |
| Tail 01 | 0.964795 | 0.968601 | 0.944814 | 0.945766 | 0.872502 | 0.870599 | 0.941009 | 0.943863 |
| Tail 02 | 0.786870 | 0.786870 | 0.734539 | 0.736441 | 0.618459 | 0.637488 | 0.757374 | 0.752617 |
| Tail 03 | 0.916270 | 0.934348 | 0.912464 | 0.929591 | 0.919125 | 0.934348 | 0.900095 | 0.901047 |
| ***COUNT*** | 1 | **7** | 0 | 1 | 0 | 1 | 0 | 0 |
| ***MEAN*** | 0.896051 | **0.904258** | 0.868577 | 0.876903 | 0.811370 | 0.821836 | 0.826832 | 0.835276 |
| ***SD*** | **0.051610** | 0.053491 | 0.067803 | 0.067431 | 0.092872 | 0.089666 | 0.075159 | 0.073106 |

### Table S8 | Comparison between BaseModel and MultiviewCorrection prediction aPCK accuracy (%) on [mouse side-view] dataset

| Data Size | FlexibleBaseline | | | DeepLabCut | | | LEAP | | | StackedDenseNet | | |
|---|---|---|---|---|---|---|---|---|---|---|---|---|
| Joint Name | BaseM | MVC | OF-MVC | BaseM | MVC | OF-MVC | BaseM | MVC | OF-MVC | BaseM | MVC | OF-MVC |
| Front Right Paw | 0.923882 | 0.932445 | 0.932445 | 0.913416 | 0.913416 | 0.912464 | 0.785918 | 0.809705 | 0.801142 | 0.735490 | 0.768792 | 0.772598 |
| Hind Right Paw | 0.888677 | 0.898192 | 0.904853 | 0.868696 | 0.871551 | 0.880114 | 0.830637 | 0.846813 | 0.843007 | 0.789724 | 0.809705 | 0.827783 |
| Front Left Paw | 0.873454 | 0.893435 | 0.895338 | 0.836346 | 0.854424 | 0.867745 | 0.761180 | 0.779258 | 0.797336 | 0.801142 | 0.823977 | 0.840152 |
| Hind Left Paw | 0.907707 | 0.910561 | 0.915319 | 0.827783 | 0.825880 | 0.835395 | 0.828735 | 0.840152 | 0.848716 | 0.793530 | 0.817317 | 0.834443 |
| Snout | 0.906755 | 0.906755 | 0.914367 | 0.910561 | 0.910561 | 0.915319 | 0.874405 | 0.874405 | 0.887726 | 0.896289 | 0.896289 | 0.902950 |
| Tail 01 | 0.964795 | 0.964795 | 0.968601 | 0.944814 | 0.944814 | 0.945766 | 0.872502 | 0.872502 | 0.870599 | 0.941009 | 0.941009 | 0.943863 |
| Tail 02 | 0.786870 | 0.786870 | 0.786870 | 0.734539 | 0.734539 | 0.736441 | 0.618459 | 0.618459 | 0.637488 | 0.757374 | 0.757374 | 0.752617 |
| Tail 03 | 0.916270 | 0.916270 | 0.934348 | 0.912464 | 0.912464 | 0.929591 | 0.919125 | 0.919125 | 0.934348 | 0.900095 | 0.900095 | 0.901047 |
| ***COUNT*** | 1 | 2 | **7** | 0 | 0 | 1 | 0 | 0 | 1 | 0 | 0 | 0 |
| ***MEAN*** | 0.896051 | 0.901165 | **0.906518** | 0.868577 | 0.870956 | 0.877854 | 0.811370 | 0.820052 | 0.827545 | 0.826832 | 0.839320 | 0.846932 |
| ***SD*** | 0.051610 | **0.051418** | 0.053296 | 0.067803 | 0.067044 | 0.067399 | 0.092872 | 0.091848 | 0.088883 | 0.075159 | 0.066080 | 0.065956 |

**Table S9 | Comparison on fully trained FlexibleBaseline model size prediction aPCK accuracy (%) on [mouse side-view] dataset**

| Joint Name | Standard | Reduced | Small |
|---|---|---|---|
| Front Right Paw | 0.923882 | 0.882969 | 0.845861 |
| Hind Right Paw | 0.888677 | 0.853473 | 0.835395 |
| Front Left Paw | 0.873454 | 0.840152 | 0.843007 |
| Hind Left Paw | 0.907707 | 0.851570 | 0.871551 |
| Snout | 0.906755 | 0.914367 | 0.921979 |
| Tail 01 | 0.964795 | 0.944814 | 0.969553 |
| Tail 02 | 0.786870 | 0.743102 | 0.679353 |
| Tail 03 | 0.916270 | 0.878211 | 0.903901 |
| ***COUNT*** | **6** | 0 | 2 |
| ***MEAN*** | **0.896051** | 0.863582 | 0.858825 |
| ***SD*** | **0.051610** | 0.059855 | 0.085874 |

Training was performed on full [mouse side-view] dataset (2599 frames), and the training was done with 50 epochs at learning rate of 0.0001, batch size of 10.

**Fig. S1 | Comparison on fully trained FlexibleBaseline model size prediction aPCK error rate (%) on [mouse side-view] dataset**

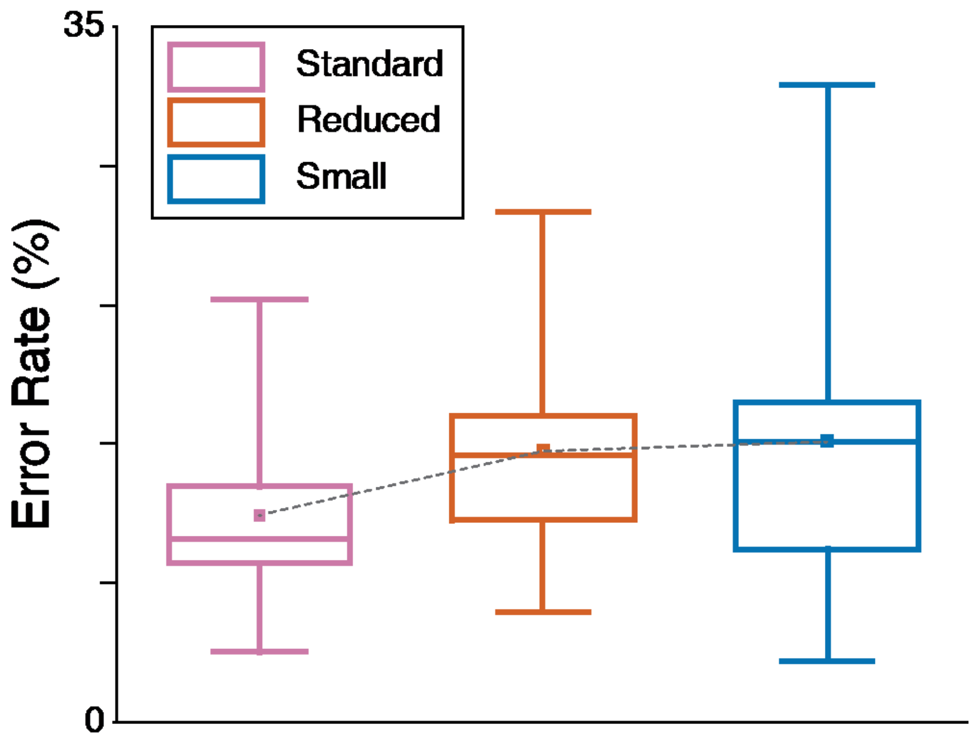


Prediction error rates represented in box plot with the five number summary being (minimum, first quartile, median, third quartile, maximum). The small square in the middle of each box plot represent the mean error rate.

**Fig. S2 | Manual labelling GUI**[1]

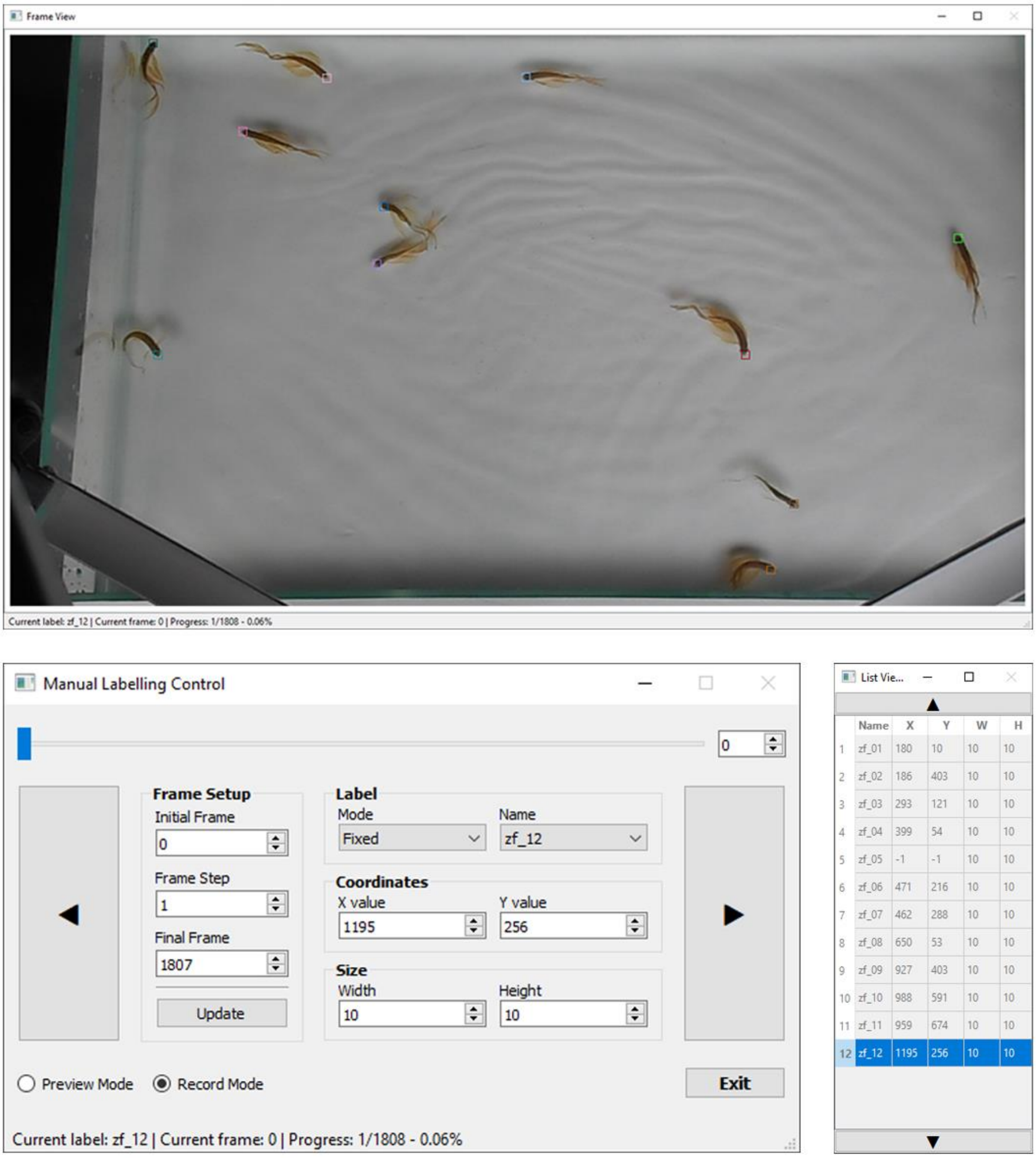


A GUI for manual labelling is included in the code package. User could use “A” and “D” key to navigate previous and next frame, “W” and “S” key to select between labels. User also able to select a label with “Left Mouse Click”, add label with “SHIFT” + “Left Mouse Click”, delete label with “Right Mouse Click”. Moving a selected label, can be achieved with simple “Mouse Dragging”.

[1] The windows in this example screenshots have been resized to fit the pages.

### Fig. S3 | Dataset creation GUI

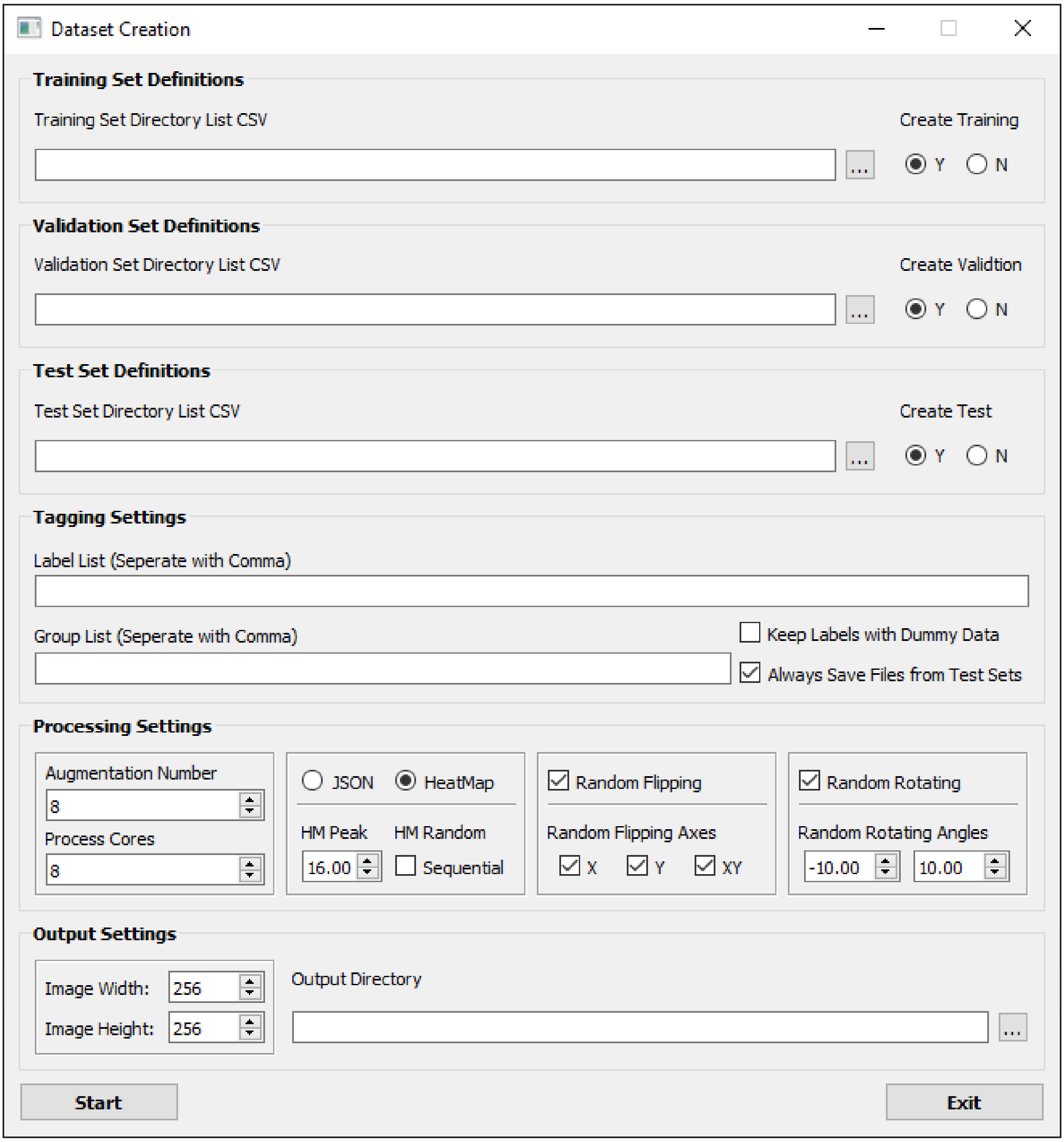


A GUI for creating learning dataset is included in the code package. This GUI could create training/validation/testing datasets form manual labelled data with many different options to meet different training requirements. All instructions have been clearly noted on the GUI.

**Fig. S4 | Dataset example**

**a**

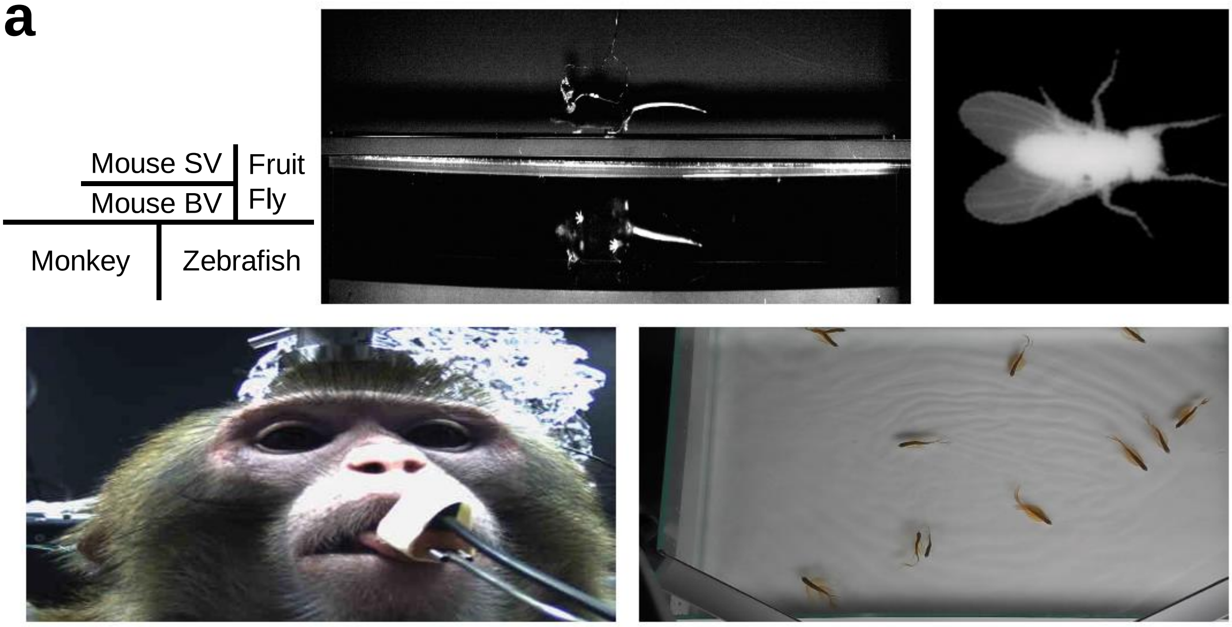


**b**

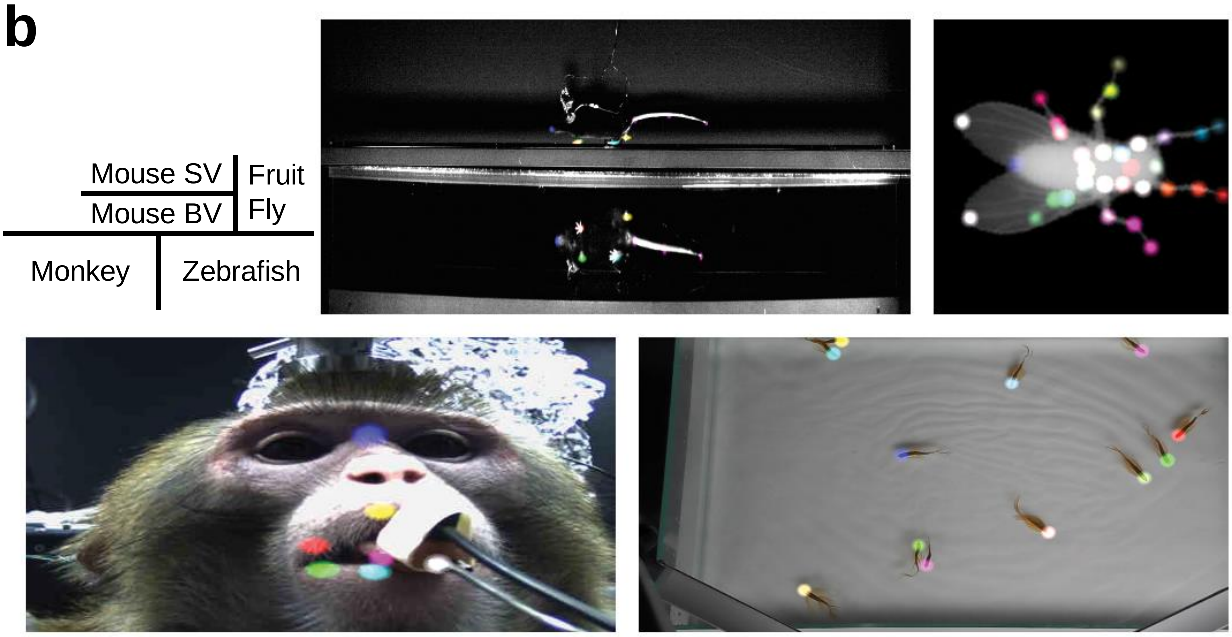


**a, Unlabelled dataset frame.**

**b, Heatmap labelled frame.**

SV = Side-view, BV = Bottom-view.
